# WGS analysis of *Listeria monocytogenes* from rural, urban, and farm environments in Norway: Genetic diversity, persistence, and relation to clinical and food isolates

**DOI:** 10.1101/2021.10.27.466212

**Authors:** Annette Fagerlund, Lene Idland, Even Heir, Trond Møretrø, Marina Aspholm, Toril Lindbäck, Solveig Langsrud

## Abstract

*Listeria monocytogenes* is a ubiquitous environmental bacterium associated with a wide variety of natural and man-made environments, such as soil, vegetation, livestock, food processing environments, and urban areas. It is also among the deadliest foodborne pathogens, and knowledge about its presence and diversity in potential sources is crucial to effectively track and control it in the food chain. Isolation of *L. monocytogenes* from various rural and urban environments showed higher prevalence in agricultural and urban developments than in forest or mountain areas, and that detection was positively associated with rainfall. Whole genome sequencing (WGS) was performed for the collected isolates and for *L. monocytogenes* from Norwegian dairy farms and slugs, in total 218 isolates. The data was compared with available datasets from clinical and food associated sources in Norway collected within the last decade. Multiple examples of clusters of isolates with 0-8 wgMLST allelic differences were collected over time in the same location, demonstrating persistence of *L. monocytogenes* in natural, urban and farm environments. Furthermore, several clusters with 6-20 wgMLST allelic differences containing isolates collected across different locations, times and habitats were identified, including nine clusters harbouring clinical isolates. The most ubiquitous clones found in soil and other natural and animal ecosystems (CC91, CC11, and CC37) were distinct from clones predominating among both clinical (CC7, CC121, CC1) and food (CC9, CC121, CC7, CC8) isolates. The analyses indicated that ST91 was more prevalent in Norway than other countries and revealed a high proportion of the hypovirulent ST121 among Norwegian clinical cases.

**Importance:** *Listeria monocytogenes* is a deadly foodborne pathogen that is widespread in the environment. For effective management, both public health authorities and food producers need reliable tools for source tracking, surveillance, and risk assessment. For this, whole genome sequencing (WGS) is regarded as the present and future gold standard. In the current study, we use WGS to show that *L. monocytogenes* can persist for months and years in natural, urban and dairy farm environments. Notably, clusters of almost identical isolates, with genetic distances within the thresholds often suggested for defining an outbreak cluster, can be collected from geographically and temporally unrelated sources. The work highlights the need for a greater knowledge of the genetic relationships between clinical isolates and isolates of *L. monocytogenes* from a wide range of environments, including natural, urban, agricultural, livestock, food production, and food processing environments, in order to correctly interpret and use results from WGS analyses.

## Introduction

*Listeria monocytogenes* is a bacterial pathogen responsible for the life-threatening disease listeriosis. The most common cause of listeriosis is considered to be ingestion of food contaminated by *L. monocytogenes* from unclean food production equipment (1, 2). *L. monocytogenes* is a ubiquitous environmental bacterium that has been associated with a wide variety of environments, such as rivers, soil, vegetation, wild and domesticated animals, food processing environments, and urban areas (3, 4). Consequently, a total absence of *L. monocytogenes* in non-heat-treated foods is difficult, perhaps impossible, to achieve. The literature is, however, not fully consistent about the main habitats of *L. monocytogenes* and the factors affecting its occurrence and spread to humans. It is therefore of importance to increase the understanding of the relationship between *L. monocytogenes* in natural and animal reservoirs, food processing environments, and human clinical disease.

The occurrence of *L. monocytogenes* in soil varies widely, from 0.7% to 45%, depending on the geographic area, season, and humidity (4-6). In comparative investigations, higher frequencies of *L. monocytogenes* have been found after rain, flooding and irrigation events (7, 8). Several studies have reported high incidence of *L. monocytogenes* in water from rivers and lakes, with frequencies from 10-62% of the samples depending on the area and detection method (9-13). A link has been found between the proximity to upstream dairy farms and cropped land and the presence of *L. monocytogenes* in river water (10, 12). An explanation for this could be high frequencies of *L. monocytogenes* in feces from farm animals, e.g., cattle, ducks, and sheep, leaking into surrounding soil and water (9, 14, 15). Dairy farms are, for example, known to hold a *L. monocytogenes* reservoir, and prevalences in environmental samples of 11-24% have been reported (6, 15-17). However, *L. monocytogenes* is not particularly linked to farm animals, and is frequently found in other animals and birds, such as game and urban birds, boars, garden slugs and rodents (9, 18-21). An association between dense populations of humans and occurrence of *L. monocytogenes* in the environment has been reported. A U.S. study showed that 4.4% of samples from urban or residential areas contained *L. monocytogenes*, while the pathogen was less frequently found in samples from forests and mountains (1.3%) (22).

*L. monocytogenes* comprises four separate deep-branching lineages, which from an evolutionary viewpoint could be considered separate species (23). These are further subdivided by multilocus sequence typing (MLST) into sequence types (STs) and clonal complexes (CCs or clones). The lineage I clones CC1, CC2, CC4, and CC6 are reported to be associated with human disease, while lineage II clones CC9 and CC121 are strongly associated with food and food processing environments (24-28). While many studies have examined the molecular genotypes of *L. monocytogenes* isolates found in food, food processing environments, and clinical disease, much less is known about the diversity present in other environments. In the few published studies, the clonal diversity in environmental samples from soil and water appears to be very high, sometimes dominated by CCs associated with disease (CC1, CC4), although other dominating clones (e.g., CC37) have also been observed (5, 13, 29). Several clones are also found in wild animals, e.g., CC7 and CC37 found in moose, boars, slugs, and game birds (18, 21, 30, 31). In environmental samples from dairy/cattle farms in Finland and Latvia, the lineage II clones CC11 (ST451), CC14, CC18, CC20, CC37, and CC91 were most predominant, while lineage I clones were rare (6, 32). Although there are some exceptions, e.g., CC1 being predominant in slugs collected in garden and farm environments in Norway (21), the majority of the clones identified in natural and farm environments do not seem to belong to CCs dominant among European food and clinical isolates.

Many studies have described persistence over time for *L. monocytogenes* clones in food processing facilities (2, 33, 34) and in individual cattle herds or farm environments (15, 35, 36). Whether *L. monocytogenes* can persist over long periods of time also in rural, urban, or agricultural environments has rarely been investigated. Studies of genetic relationships between *L. monocytogenes* isolates from natural and animal reservoirs and isolates from food and clinical sources are scarce. High resolution molecular fingerprinting based on whole genome sequencing (WGS) technology has revolutionized the ability to detect outbreaks and the presence of persistent strains (37). Few studies have, however, carried out WGS analyses of *L. monocytogenes* isolates collected from non-food associated locations over the span of months and years. The present study aimed to use WGS to investigate the diversity and genetic relationships between *L. monocytogenes* isolates from rural, agricultural, and urban environments in Norway, and to compare these with available datasets containing genomes of *L. monocytogenes* from human clinical and food-associated sources in Norway collected within the last decade.

## Results

### Prevalence of *L. monocytogenes* in rural and urban environments in Norway

A total of 618 distinct environmental sites from rural and urban environments were sampled for *L. monocytogenes* between April 2016 and April 2020. The overall sampling scheme was designed to obtain an overview of the presence of *L. monocytogenes* in various habitats, and samples were collected from several geographical regions in Norway (Figure 1A). To study potential persistence of *L. monocytogenes* clones over time, some sites were sampled more than once. At the onset of the study, we hypothesized that the presence of *L. monocytogenes* would be more strongly associated with farm animals, agricultural activity, and urban areas than with natural forests and other wildlands (22). During the first sampling occasion, 10% of sample sites were positive for *L. monocytogenes* (Table 1). In addition, 13 samples of commercial bags of plant soil or compost were negative for *L. monocytogenes*. In concordance with our hypothesis, the prevalence of *L. monocytogenes* was significantly higher in urban areas and in areas associated with agriculture and livestock (agricultural fields, grazelands and animal paths) than in forest/mountain areas and on footpaths (Fisher’s exact test *p*<0.02). Sampling locations classified as footpaths were generally from non-urban areas such as woods or other areas used for hiking. While 14% of urban areas were positive for *L. monocytogenes*, all samples from footpaths were negative for *L. monocytogenes*, and only 2% of the samples collected in woodland or mountain areas were positive.

**Table 1:** Prevalence of L. monocytogenes in rural and urban environments

| Habitat or sampling area | No. of collected isolates | Prevalence of *L. monocytogenes* (%) |
| --- | --- | --- |
| Grazeland or animal path | 85 | 21 % |
| Urban or residential area | 177 | 14 % |
| Agricultural field | 70 | 11 % |
| Near food processing plant | 106 | 10 % |
| Beach or sandbank | 24 | 4 % |
| Forest or mountain area | 121 | 2 % |
| Footpaths | 35 | 0 % |

**Figure 1:**
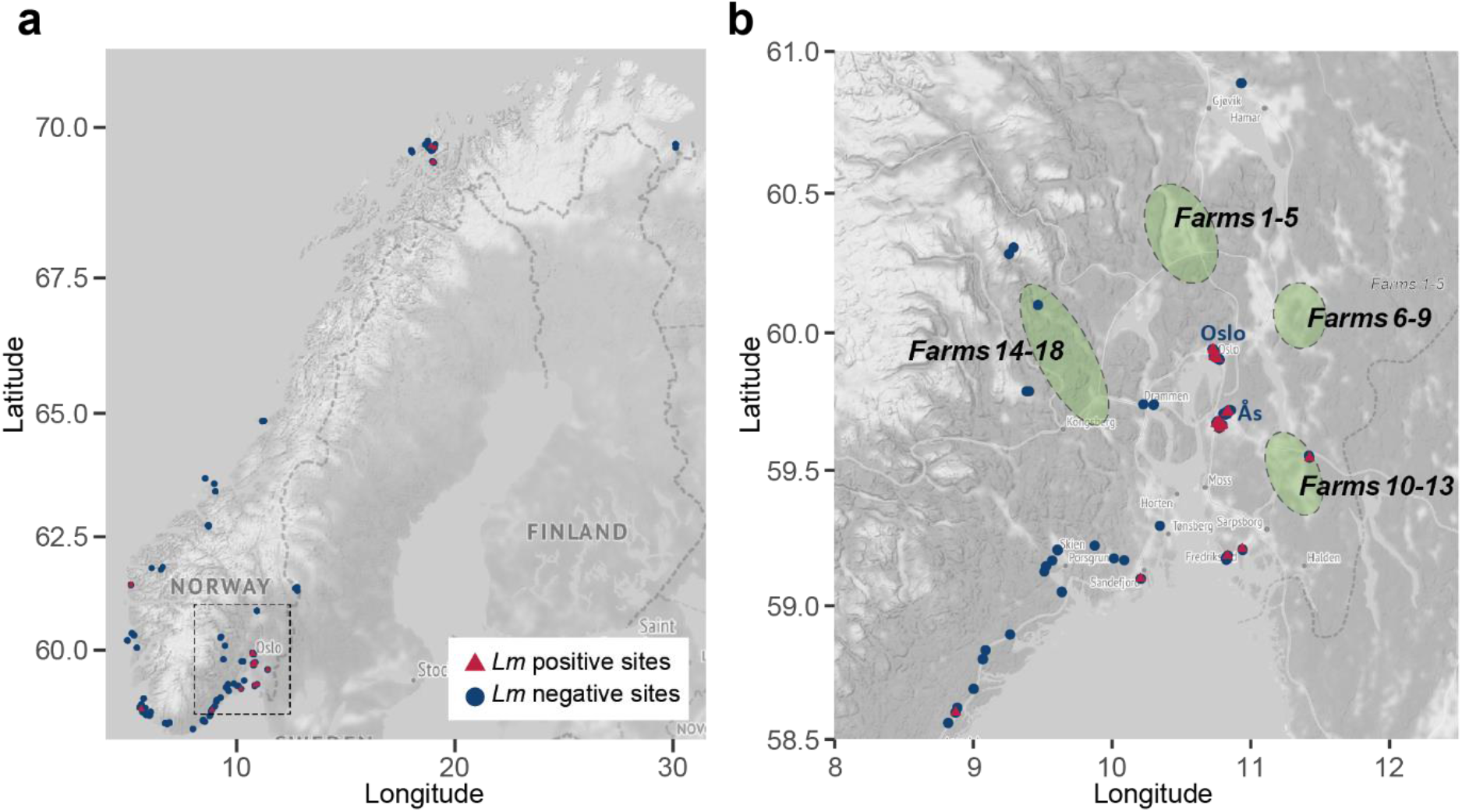
Maps showing the geographic location of sampling sites. **A)** The location of sampling sites in rural and urban environments in Norway, with red triangles representing *L. monocytogenes* positive samples and blue circles negative sampling points. The area outlined by the dashed square in **a)** is the area shown in **b)**. The green shaded areas in **b)** show the geographical origins of the dairy cattle farms sampled for *L. monocytogenes* in Idland *et al*. (16).

### Detection of *L. monocytogenes* correlated with rain and sample humidity

Previous studies have indicated that *L. monocytogenes* is more frequently isolated after recent rainfall, irrigation, and flooding events (38, 39). In the present study, 271 out of 618 samples were collected on days with rainfall and 347 samples on days with no rain within the previous 24 hours. When collected on days with rain, 20% of samples were positive for *L. monocytogenes*, while on days with no rain within the last 24 hours, only 3% of samples were positive. Thus, our data supports previous studies suggesting that prevalence of *L. monocytogenes* is positively associated with rainfall (Fisher’s exact test *p*=2×10^−12^).

Upon sample collection, the humidity of the sampled material was categorized on a scale from 1 (completely dry) to 5 (liquid). Overall, the prevalence of *L. monocytogenes* in samples from the two driest sample categories was 5.6% (4/70) and 5.7% (11/196), while it was 17% (27/164) and 14% (12/86) in the more humid categories 3 and 4. The prevalence was significantly higher in the humid samples (categories 3 and 4) than in the two driest sample categories (Fishers exact test, *p*<0.02). The prevalence in liquid samples (category 5) was 10% (10/102). Among the samples collected in urban environments, the sample humidity was not significantly associated (*p*>0.05) with the prevalence of *L. monocytogenes*, with an overall prevalence of 10% in categories 1 and 2 (10/92), and 17% in categories 3 to 5 (14/85).

### Persistent strains detected in rural and urban environments

To examine whether environmental locations retained their status as *L. monocytogenes* positive or negative over time, and whether the same clones were isolated repeatedly from the same location, 70 sites were subjected to one to three additional rounds of sampling the following years. In total, 115 *L. monocytogenes* isolates were collected in the current study (S1 Table). All isolates were subjected to WGS, *in silico* MLST, and whole genome MLST (wgMLST) analysis. The distribution of clones (CCs) among the identified isolates is presented in Figure 2.

**Figure 2:**
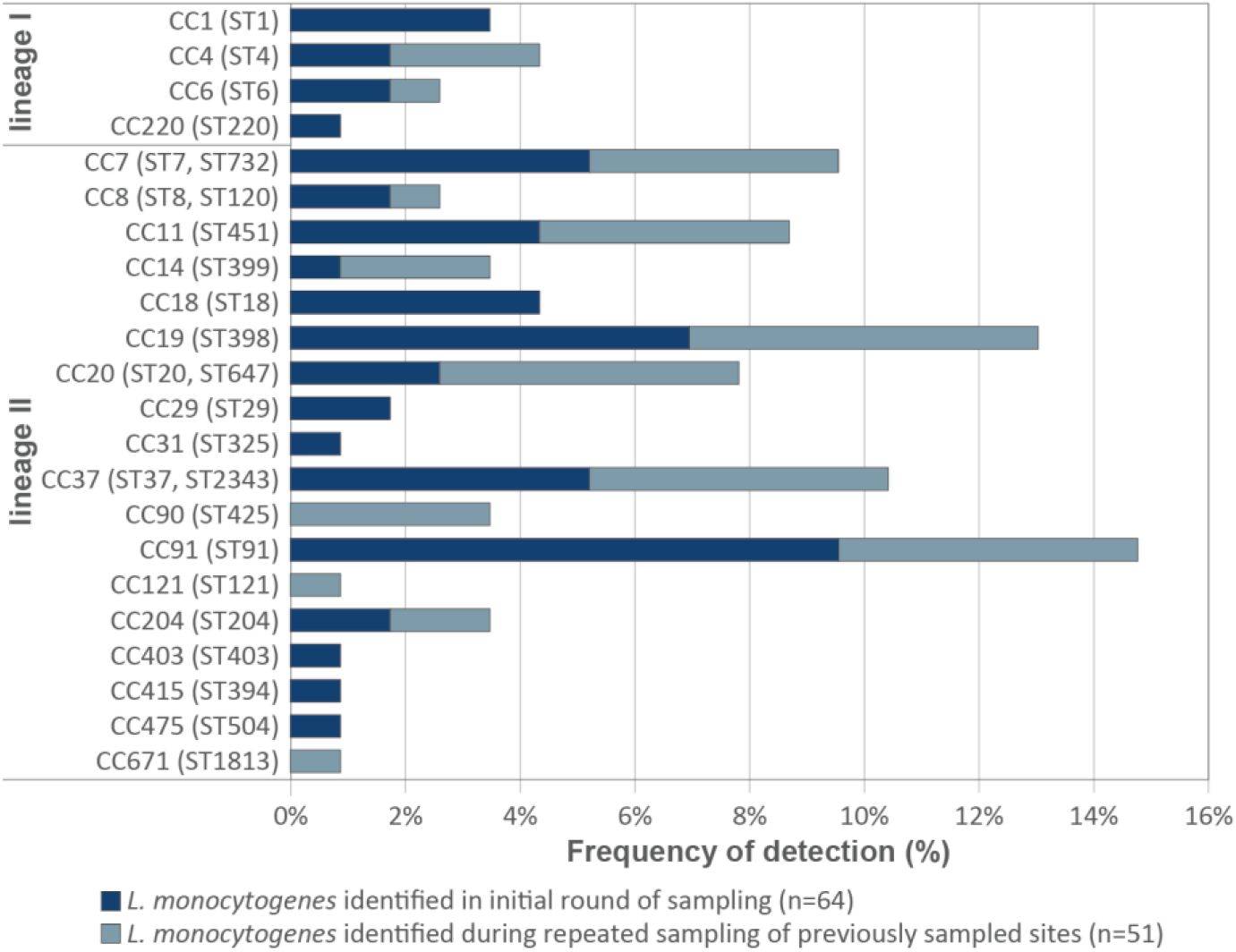
Distribution of CCs among identified isolates from rural and urban environments. The data is reported as percentage of the grand total number of isolates (n=115). STs represented within each clonal complex (CC) are given in parentheses.

Of the 44 sampling points positive for *L. monocytogenes* in the first round of sampling, 28 sites (64%) were positive on at least one of the subsequent sampling occasions. Of the 26 initially negative sites, five turned out positive during later sampling events (19%), and one of these was positive twice. In total, 29 sampling sites were positive for *L. monocytogenes* more than once, and isolates belonging to the same ST was collected repeatedly from seven sites (Table 2). In six cases, STs repeatedly isolated from the same site were very closely related, with a maximum wgMLST allelic distance of 20. When also adjacent or slightly more distant sampling sites (max 3 km) were included, a total of 14 clusters with genetic distances <20 were repeatedly collected from the same location over periods ranging from 4 months to 3 years (S2 Table and S1 Text). When the commonly employed core genome MLST (cgMLST) scheme described by Moura *et al*. (23) was employed, the isolates could not be distinguished, except in one cluster with distances of 0 to 1 cgMLST alleles. Twelve clusters, including two clusters each for CC91, CC11 (ST451) and CC37, represent clusters of highly similar isolates with range 0 to 8 wgMLST allelic differences. Together, the results strongly indicate that *L. monocytogenes* clones had persisted in the same environment or were repeatedly reintroduced between sampling events in both rural and urban locations.

**Table 2:** STs identified at sampling points positive for L. monocytogenes on repeated occasions

| Site no. | Sampling point description | 2016, Oct | 2017, Jun, Oct, Nov | 2018, Jun | 2018, Sept, Oct | 2019, Sept | 2020, Jan |
| --- | --- | --- | --- | --- | --- | --- | --- |
| **Urban or residential area** |  |  |  |  |  |  |  |
| 33 | Brook in residential area | ST451 |  |  |  | ST4 |  |
| 48 | Garden compost heap | ST451<sup>a</sup> |  |  |  | ST451<sup>a</sup> | ST425 |
| 49 | Garden compost heap | ST451 |  |  |  | ST425<sup>b</sup> | ST425<sup>b</sup> |
| 66 | Puddle next to road | ST20 |  |  |  | ST4 |  |
| 120 | Flowerbed in town centre |  | ST91 |  | negative | ST1813 | ST398 |
| 121 | In front of park bench by flowerbed |  | ST399 |  | ST451 | ST398<sup>c</sup> | ST398<sup>c</sup> |
| 123 | Grass lawn in town centre |  | ST398<sup>d</sup> |  | ST398<sup>d</sup> | negative | negative |
| 129 | Roadside close to brook |  | ST37 |  |  | ST91 | negative |
| 251 | Decaying leaves/vegetation on bike path |  | ST18 |  |  | ST37<sup>e</sup> | ST37<sup>e</sup> |
| 252 | Soil near horse paddock |  | ST204 |  |  | ST398 | ST7 |
| 253 | Along sidewalk curb |  | ST1 |  |  | negative | ST425 |
| 259 | Flowerbed with pigeons, city centre |  | ST204<sup>f</sup> |  |  | ST204<sup>f</sup> | ST398 |
| 262 | At foot of tree, city centre |  | ST6 |  |  | ST204 | ST120 |
| 268 | Puddle on gravel path, city park |  | ST6 |  |  | ST451 | negative |
| **Grazeland or animal path** |  |  |  |  |  |  |  |
| 53 | Decaying vegetation by feeding station | ST4 |  |  |  | ST399 | ST37 |
| 54 | Soil close to cattle feeding station | ST91 |  |  |  | ST399 | ST451 |
| 55 | Mud close to cattle enclosure | ST37 |  |  |  | ST399 | ST91 |
| 56 | Puddle of mud close to cattle enclosure | ST91 |  |  |  | ST398 | ST6 |
| 98 | Sheep grazing pasture |  | ST398 |  | ST451 | ST2343 | ST91 |
| 99 | Sheep manure |  | ST398 |  | negative | negative | ST91 |
| 100 | Animal tracks by feeding station |  | ST398 |  | negative | ST37 | negative |
| 101 | Animal tracks by feeding station |  | ST398 |  | negative | ST91 | negative |
| 130 | Soil at edge of pond |  | ST91 |  | negative | ST121 | negative |
| 133 | Decaying vegetation at edge of pond |  | ST398 |  | negative | ST20<sup>g</sup> | negative |
| 134 | Decaying vegetation at edge of pond |  | ST29 |  | negative | ST4 | ST20<sup>g</sup> |
| **Near food processing plant** |  |  |  |  |  |  |  |
| 279 | Grass next to cold storage entrance |  | ST732 | ST732 | ST732 |  |  |
| 287 | Storm drain outside plant |  | ST1 | negative | ST647 |  | negative |
| 363 | Gravel from quay outside factory |  |  | negative | ST732 |  | ST647 |
| 406 | Gravel from quay outside factory |  |  |  | ST1 |  | ST647 |

Allelic distance between isolates with same superscript letter: <sup>a</sup> 3 wgMLST differences, <sup>b</sup> 2 wgMLST differences, <sup>c</sup> 2 wgMLST differences, <sup>d</sup> 34 wgMLST differences and 0 cgMLST differences, <sup>e</sup> 7 wgMLST differences, <sup>f</sup> 3 wgMLST differences, <sup>g</sup> 0 wgMLST differences (sites 133 and 134 are located 5 m apart), <sup>h</sup> 0-4 wgMLST differences.

We also observed a case where a recent common contamination source was obscure: only 9 wgMLST alleles (and no cgMLST alleles) separated a pair of CC6 isolates found 30 km and 3 years apart; one isolate from a grazing pasture in Akershus county in 2020 and the other from soil by the root of a tree in Oslo city centre in 2017.

### Persistence and cross-contamination on Norwegian dairy farms

In the next step, WGS was performed for a panel of 79 *L. monocytogenes* isolates collected from Norwegian dairy farms (16). A total of 18 dairy herds from four different geographical areas within a 100 km radius from downtown Oslo (Figure 1B) had each been sampled four to six times between August 2019 and July 2020. Out of the 556 analyzed samples, *L. monocytogenes* was detected in 12 milk filters (13% prevalence), 30 cattle feces samples (30%), 32 samples of cattle feed (silage or silage mixture; 32%), and 5 teat swabs (5%). All bulk tank milk and teat milk samples were negative for *L. monocytogenes*, and for one of the farms (farm 16), all 34 collected samples were negative. An overview of the STs of the collected isolates (S1 Table) is presented in Table 3, and a phylogenetic tree showing the genetic relationships between the individual isolates is shown in Figure 3.

**Table 3:**
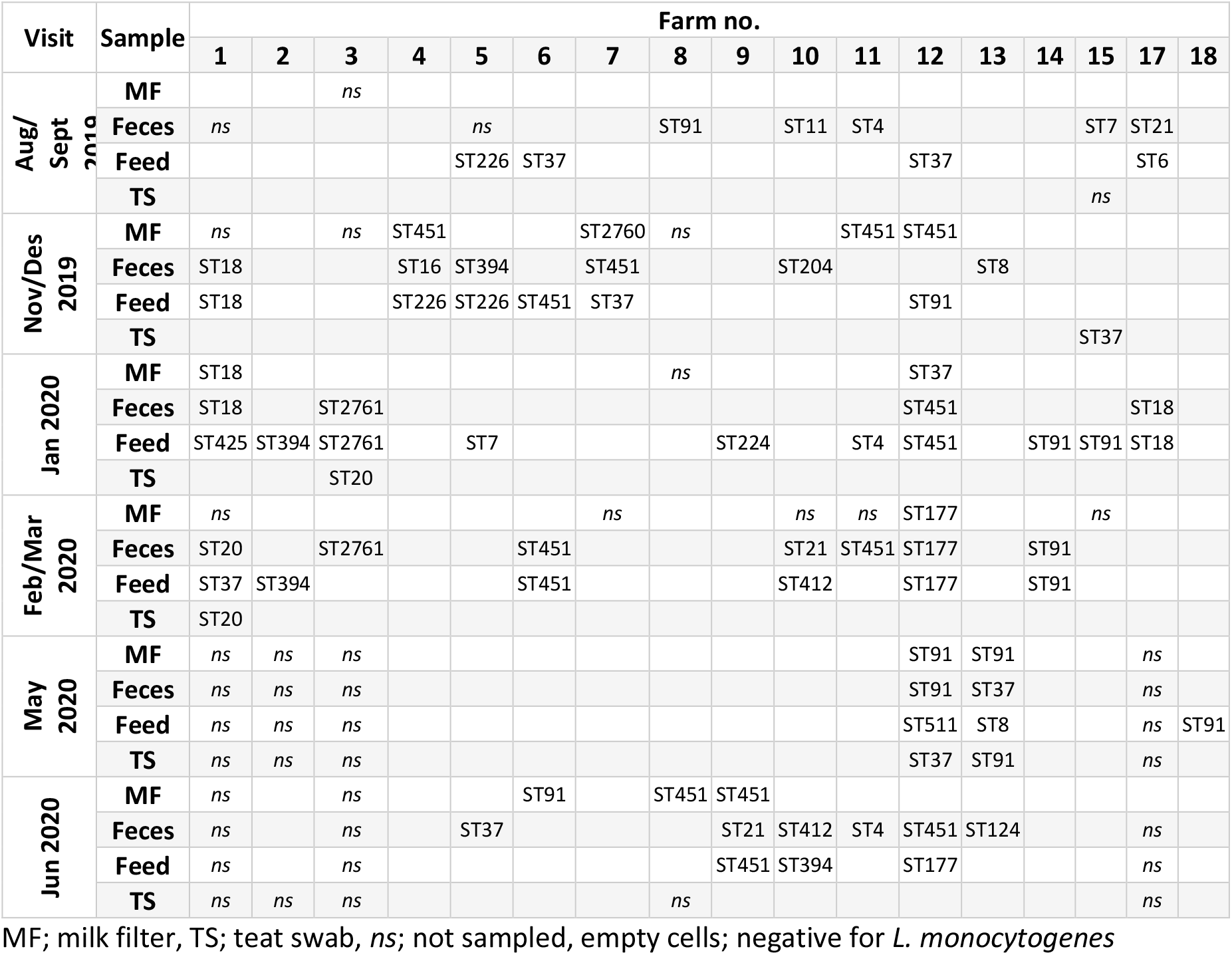
L. monocytogenes STs identified on dairy farms

| Visit | Sample | Farm no. |  |  |  |  |  |  |  |  |  |  |  |  |  |  |  |  |
| --- | --- | --- | --- | --- | --- | --- | --- | --- | --- | --- | --- | --- | --- | --- | --- | --- | --- | --- |
|  |  | 1 | 2 | 3 | 4 | 5 | 6 | 7 | 8 | 9 | 10 | 11 | 12 | 13 | 14 | 15 | 17 | 18 |
| Aug/<br>Sept<br>2019 | MF |  |  | ns |  |  |  |  |  |  |  |  |  |  |  |  |  |  |
|  | Feces | ns |  |  |  | ns |  |  | ST91 |  | ST11 | ST4 |  |  |  | ST7 | ST21 |  |
|  | Feed |  |  |  |  | ST226 | ST37 |  |  |  |  |  | ST37 |  |  |  | ST6 |  |
|  | TS |  |  |  |  |  |  |  |  |  |  |  |  |  |  | ns |  |  |
| Nov/Dec<br>2019 | MF | ns |  | ns | ST451 |  |  | ST2760 | ns |  |  | ST451 | ST451 |  |  |  |  |  |
|  | Feces | ST18 |  |  | ST16 | ST394 |  | ST451 |  |  | ST204 |  |  | ST8 |  |  |  |  |
|  | Feed | ST18 |  |  | ST226 | ST226 | ST451 | ST37 |  |  |  |  | ST91 |  |  |  |  |  |
|  | TS |  |  |  |  |  |  |  |  |  |  |  |  |  |  | ST37 |  |  |
| Jan 2020 | MF | ST18 |  |  |  |  |  |  | ns |  |  |  | ST37 |  |  |  |  |  |
|  | Feces | ST18 |  | ST2761 |  |  |  |  |  |  |  |  | ST451 |  |  |  | ST18 |  |
|  | Feed | ST425 | ST394 | ST2761 |  | ST7 |  |  |  | ST224 |  | ST4 | ST451 |  | ST91 | ST91 | ST18 |  |
|  | TS |  |  | ST20 |  |  |  |  |  |  |  |  |  |  |  |  |  |  |
| Feb/Mar<br>2020 | MF | ns |  |  |  |  |  | ns |  |  | ns | ns | ST177 |  |  | ns |  |  |
|  | Feces | ST20 |  | ST2761 |  |  | ST451 |  |  |  | ST21 | ST451 | ST177 |  | ST91 |  |  |  |
|  | Feed | ST37 | ST394 |  |  |  | ST451 |  |  |  | ST412 |  | ST177 |  | ST91 |  |  |  |
|  | TS | ST20 |  |  |  |  |  |  |  |  |  |  |  |  |  |  |  |  |
| May<br>2020 | MF | ns | ns | ns |  |  |  |  |  |  |  |  | ST91 | ST91 |  |  | ns |  |
|  | Feces | ns | ns | ns |  |  |  |  |  |  |  |  | ST91 | ST37 |  |  | ns |  |
|  | Feed | ns | ns | ns |  |  |  |  |  |  |  |  | ST511 | ST8 |  |  | ns | ST91 |
|  | TS | ns | ns | ns |  |  |  |  |  |  |  |  | ST37 | ST91 |  |  | ns |  |
| Jun 2020 | MF | ns |  | ns |  |  | ST91 |  | ST451 | ST451 |  |  |  |  |  |  |  |  |
|  | Feces | ns |  | ns |  | ST37 |  |  |  | ST21 | ST412 | ST4 | ST451 | ST124 |  |  | ns |  |
|  | Feed | ns |  | ns |  |  |  |  |  | ST451 | ST394 |  | ST177 |  |  |  | ns |  |
|  | TS | ns | ns | ns |  |  |  |  | ns |  |  |  |  |  |  |  | ns |  |

MF; milk filter, TS; teat swab, ns; not sampled, empty cells; negative for *L. monocytogenes*

**Figure 3:**
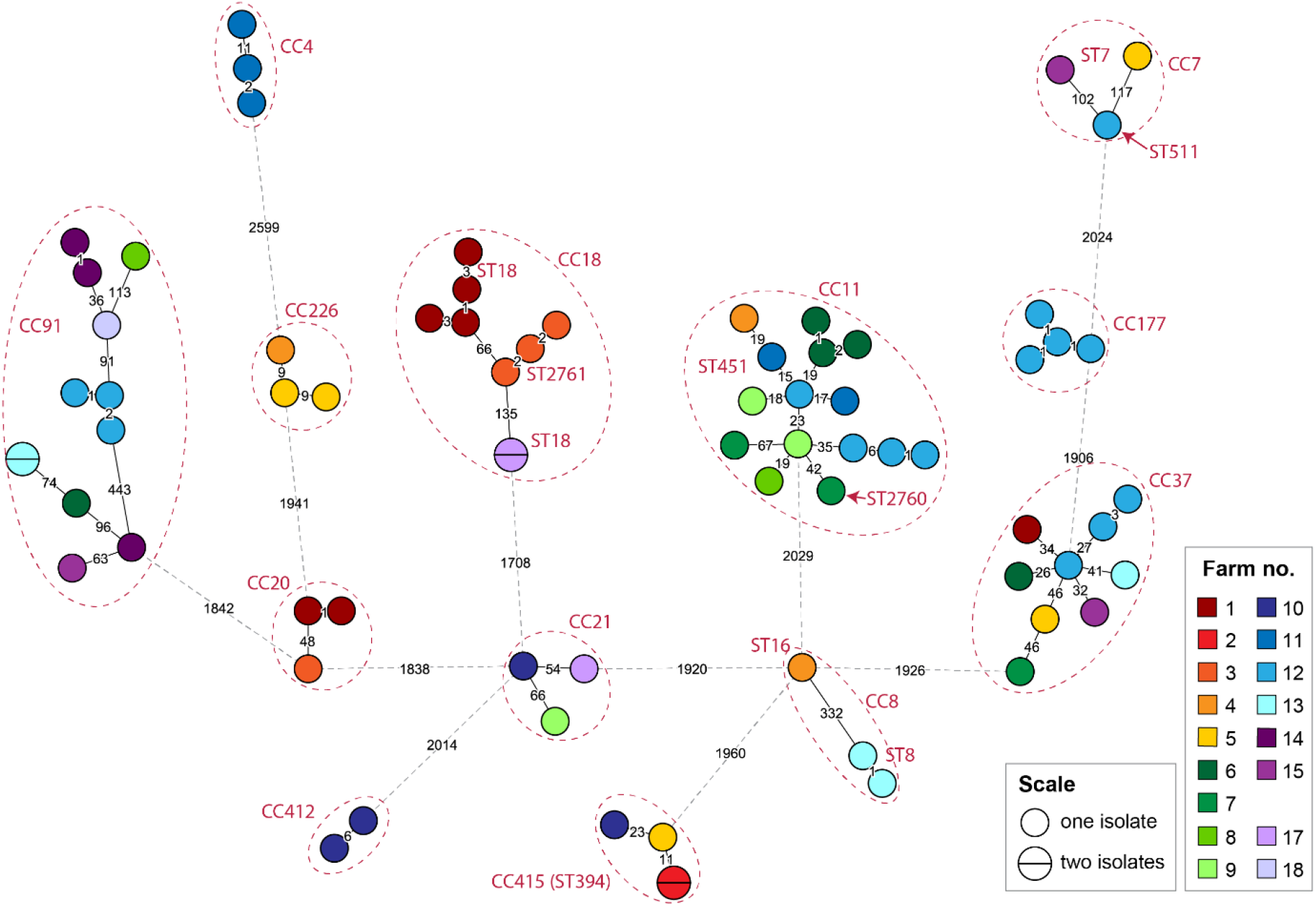
Phylogeny for the *L. monocytogenes* isolates from dairy farms. A minimum spanning tree based on wgMLST analysis. The area of each circle is proportional to the number of isolates represented, and the number of allelic differences between isolates is indicated on the edges connecting two nodes. The CCs and STs are indicated next to each node (the CC number is the same as the ST number unless indicated). Edges shown as dashed lines separate clusters belonging to different clonal complexes. Isolates separated from the nearest other isolate by >1700 wgMLST alleles (D011L, D058L, D080L, D084L, D144L, and D190L) were excluded from the figure for clarity.

Twelve clusters, each comprising two to four isolates, with pairwise genetic distances in the range 0 to 11 wgMLST alleles, were isolated from the same farm during repeated multiple visits, over periods ranging from 2 to 10 months. These clusters involved 33 of the collected isolates and comprised ten different CCs (S3 Table). These observations strongly support previous studies indicating that the same *L. monocytogenes* clones can persist over time in individual cattle herds or farm environments (15, 35, 36).

Out of 12 isolates from milk filters, four belonged to a persistent cluster, and one was closely related to an isolate from a teat swab sample obtained on the same sampling occasion. When the same clone was isolated from several sampling sites at the same farm, the pairwise genetic distances separating milk filter isolates from fecal, feed or teat swab isolates ranged from 0 to 7 wgMLST allelic differences (S3 Table). These links represent likely cross-contamination events where milk filters (and consequently milk) have been contaminated with *L. monocytogenes* clones found in the farm environment.

### Detection of closely related isolates from different geographic areas

In four cases, closely related isolates belonging to CC11 (ST451), CC226, and CC415 (ST394), were collected from more than one dairy farm. The genetic differences between isolates from different farms were somewhat greater than the diversity between isolates found on the same farm, with between 9 and 20 pairwise wgMLST allelic differences. The number of cgMLST differences within each cluster was 0 or 1 (S3 Table and S1 Text). These data indicate that farms located at different geographical areas may host the same strain of *L. monocytogenes*.

Six clusters comprising *L. monocytogenes* from both dairy farms and isolates obtained from rural and urban environments were detected. The genetic distances separating isolates from the two datasets in these clusters ranged from 9 to 27 wgMLST allelic differences, and 0 or 1 cgMLST differences (S4 Table and S1 Text). The closest link was observed for a cluster of four CC37 isolates; two from grazing land/pasture in the vicinity of Ås, and two from feed and teat swab samples obtained on two different visits to farm 12, located about 50 km east of Ås. The two pairs of isolates were separated by 9 to 14 wgMLST allelic differences and indistinguishable by cgMLST.

To further explore the occurrence of genetic links between Norwegian isolates from natural and animal reservoirs, 24 of the 34 *L. monocytogenes* isolates collected from invading slugs (*Arion vulgaris*) from garden and farm environments in Norway by Gismervik *et al*. in 2012 (21) were subjected to WGS analysis (S1 Table). Interestingly, two pairs of slug isolates collected from different geographic locations differed by only 2 (CC14) and 11 (CC1) wgMLST allelic differences. Furthermore, five clusters with 10 to 21 wgMLST allelic differences comprised a slug isolate and one or more isolates from either a rural/urban sampling site or from a dairy farm (S5 Table). The closest genetic relationship concerned two CC1 isolates, in which a slug isolate from the west coast of Norway (collected in 2012) showed only 10 wgMLST allelic differences compared to an isolate collected from a street in a residential area in Oslo in 2017.

Thus, counting the previously mentioned pair of CC6 isolates collected in Akershus and Oslo, a total of 17 close genetic links between isolates collected at relatively distant geographic areas in Norway were detected in the set of 218 examined isolates. Presumably, not all clusters represent direct epidemiological links, especially in cases where isolates were collected several years apart. The observed genetic distances within the clusters; ≤21 wgMLST and ≤3 cgMLST allelic differences, are within the thresholds often suggested as an appropriate guide for defining an outbreak cluster, which is about 7-10 cgMLST differences (23, 40, 41) or about 20 single nucleotide polymorphisms (SNPs) in SNP analyses (42, 43), which have a sensitivity comparable to wgMLST (34, 44).

### Comparison with Norwegian clinical isolates

The identification of close genetic links between isolates from different natural and animal-associated sources without known connections led us to hypothesise that it would be possible to identify clusters containing both environmental and clinical isolates with a similar level of genetic relatedness. A dataset of Norwegian clinical isolates was identified, comprising 130 genomes from 2010-2015 (92% of all reported cases in these years) (45) and two genomes from 2018 (ST20 and ST37), made publicly available by the European Centre for Disease Prevention and Control (ECDC) and the Norwegian Institute of Public Health (NIPH), respectively. Sequencing data of sufficient quality for wgMLST analysis was available for 111 of these isolates (S1 Table). An initial comparison between the clinical isolates identified 15 pairs of isolates and nine larger clusters comprising 3 to 12 isolates showing genetic distances of ≤10 cgMLST allelic differences (S6 Table). Most clusters contained isolates collected during a time span of several years and could potentially represent listeriosis outbreaks or epidemiologically linked cases.

A wgMLST analysis showing the genetic relationships between isolates originating from rural and urban environments, dairy farms, slugs, and clinical cases is shown in Figures 4 and 5. Nine clusters contained clinical isolates differentiated from isolates sequenced in the current study by genetic distances in the range 6 to 23 wgMLST allelic differences (0 to 7 cgMLST alleles) (S7 Table and S1 Text). The environmental *L. monocytogenes* closely related to clinical isolates comprised isolates from soil samples from both urban and rural locations (belonging to CC4, CC7, CC11/ST451, CC220, CC403, and CC415/ST394), three slug isolates obtained from garden and farm environments (CC7, CC8, and CC9) and a group of CC11/ST451 isolates from dairy farms. The closest genetic link was found between the single CC9 slug isolate (from 2012) and a clinical isolate from 2015, differentiated by only 6 wgMLST alleles (and 0 cgMLST alleles). The analysis shows that *L. monocytogenes* isolates that are genetically very closely related to clinical isolates can be detected in various natural and agricultural environments, even when isolates are collected across timespans ranging several years.

**Figure 4:**
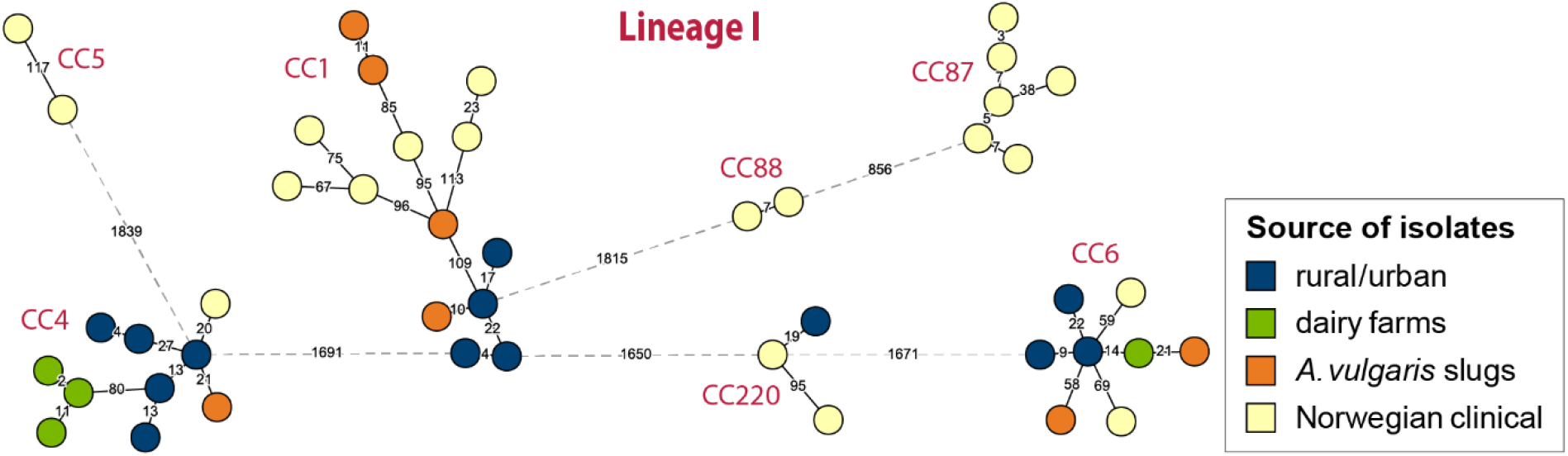
wgMLST phylogeny for *L. monocytogenes* lineage I isolates from Norway. Shown is a minimum spanning tree based on wgMLST analysis. The area of each circle is proportional to the number of isolates represented, and the number of allelic differences between isolates is indicated on the edges connecting two nodes. Edges shown as dashed lines separate clusters belonging to different clonal complexes, which are indicated next to each node. Lineage I isolates separated from the nearest other isolate by >900 wgMLST alleles (D084L, ERR2522285, ERR2522267, and ERR2522291) were excluded from the figure for clarity.

**Figure 5:**
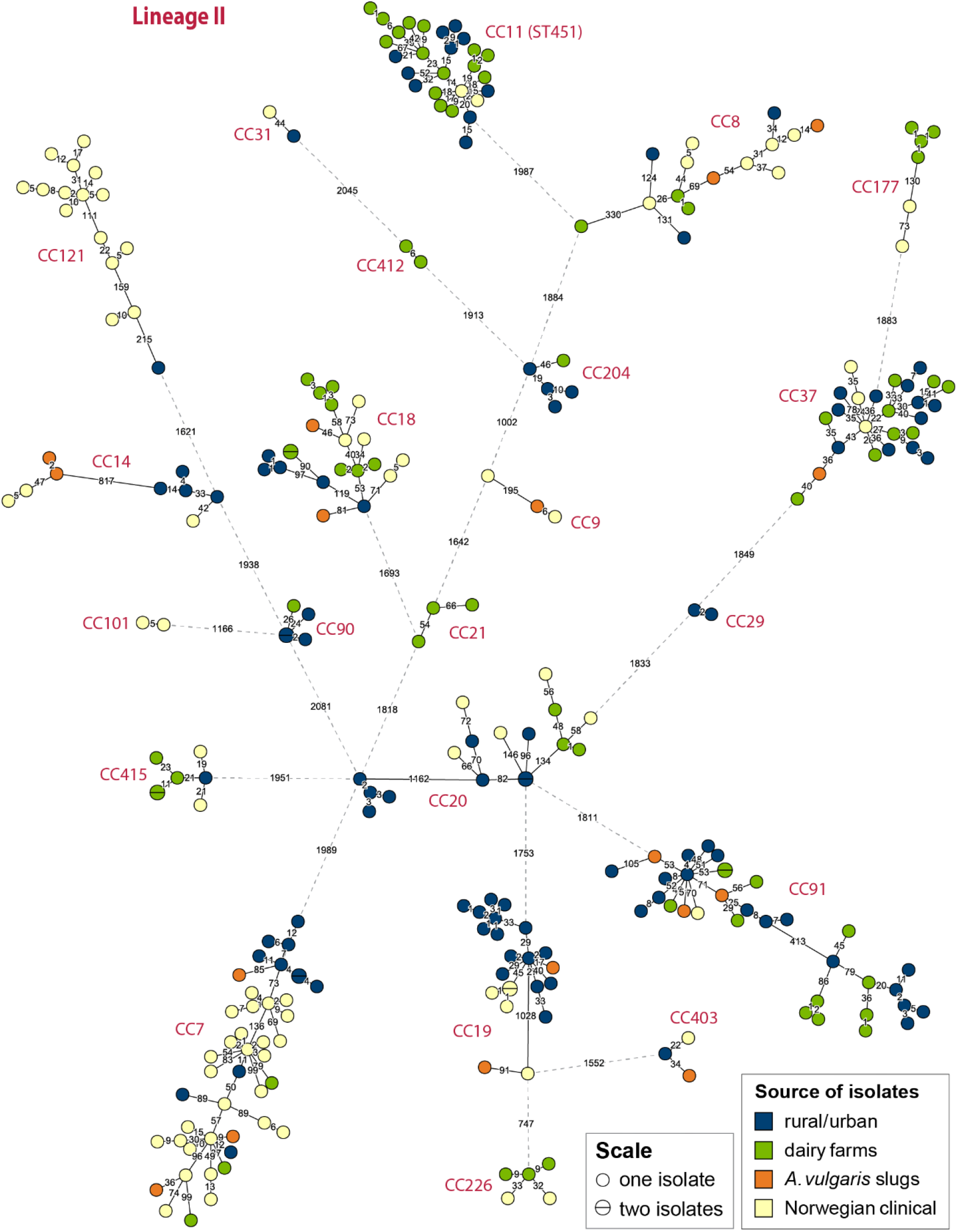
wgMLST phylogeny for *L. monocytogenes* lineage II isolates from Norway. Shown is a minimum spanning tree based on wgMLST analysis. The area of each circle is proportional to the number of isolates represented, and the number of allelic differences between isolates is indicated on the edges connecting two nodes. Edges shown as dashed lines separate clusters belonging to different clonal complexes, which are indicated next to each node. Lineage II isolates separated from the nearest other isolate by >900 wgMLST alleles (MF7614, MF6841, D144L, D190L, ERR2522251, and ERR2522298) were excluded from the figure for clarity.

### Comparison of prevalence and diversity of MLST clones from different sources

Most isolates from natural and agricultural environments belonged to *L. monocytogenes* lineage II, comprising 89%, 94%, and 68% of isolates from rural/urban environments, dairy farms, and slugs, respectively (Figure 6A). The remaining isolates belonged to *L. monocytogenes* lineage I, as lineage III or IV isolates were not detected in the current study. The predominant clones among the rural/urban isolates were CC91 (15%), CC19/ST398 (13%), CC37 (10%), and CC11/ST451 (9%). No specific niches were found for these clones as isolates were spread geographically (3-5 counties), found in 3-5 different habitats/areas and in a range of humidity and weather conditions. CC91 appeared most ubiquitous, as it was isolated from five different counties, from different sample types (soil, sand, vegetation, and feces), from five different areas (agricultural fields, urban area, beach, grazeland and forest), during all seasons and from all categories of humidity. CC11/ST451, CC91, and CC37 were also the most frequently isolated clonal groups at the dairy farms (18%, 15%, and 11%, respectively); each detected on seven different farms. Among the slug isolates, the most common clones were CC1 (15) and CC91 (12%) (21). A survey of previous studies indicated that CC1, CC7 and CC37 were the clones most commonly detected in various natural and farm environments (S8 Table).

**Figure 6:**
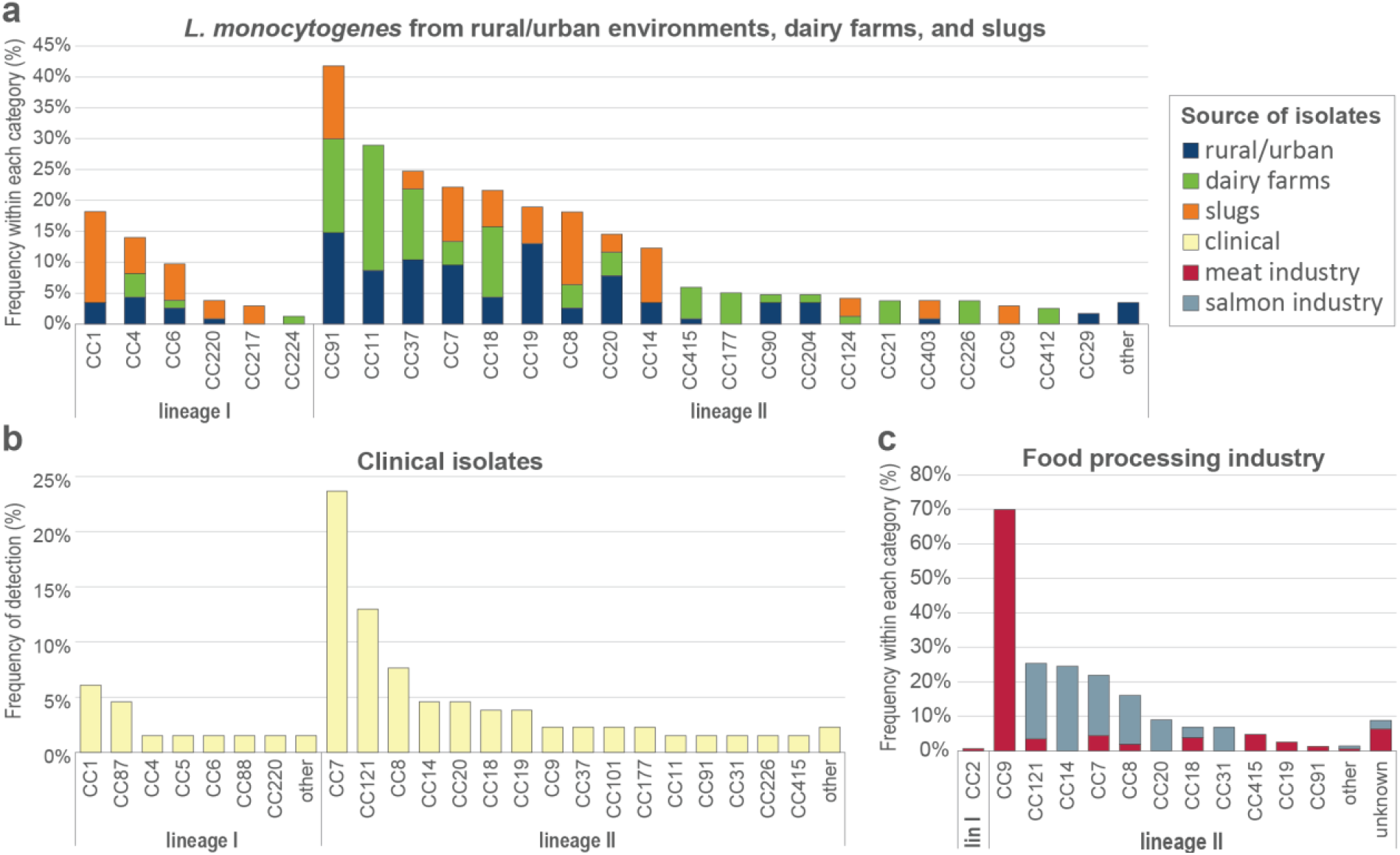
Prevalence and distribution of *L. monocytogenes* MLST clonal complexes (CCs) from different sample types in Norway. The data are reported as percentages of isolates within a given CC in each source category. **a)** Prevalence in rural and urban environments (isolated during 2016-2020; n=115 isolates), dairy farms (2019-2020; n=87), and slugs (2012; n=34). The “other” category comprises one isolate each for CC31, CC121, CC475, and CC671. **b)** Prevalence in publicly available genomes from human cases of listeriosis in Norway (2010-2015; n=129 and 2018; n=2). The lineage I “other” category comprises a CC3 and a CC59 isolate, and the lineage II “other” category includes one isolate each for CC11, CC101, and CC177. **c)** Prevalence within food processing facilities in Norway. The CCs were inferred for isolates from five meat and four salmon processing facilities (meat: 2012-2015; n=293, and salmon: 2011-2014; n=358). The data used to predict the CC for each isolate were multiple-locus variable-number tandem repeat analysis (MLVA) obtained for all isolates and MLST data obtained for representative isolates from each obtained MLVA profile (47). The “unknown” category represents isolates with MLVA profiles identified only once and not subjected to MLST.

Among the examined Norwegian clinical isolates, CC7 was the most prevalent clonal group, accounting for 23% (n=30) of the reported listeriosis cases, followed by CC121 (13%), CC8 (8%), and CC1 (6%) (Figure 6B). In contrast to that observed in many other countries (27, 46), lineage I isolates comprised a minority of the clinical isolates in this dataset (20%). The high prevalence of CC121 among the clinical isolates was unexpected, as this clone is commonly regarded as hypovirulent due to the frequent occurrence of premature stop codons (PMSC) in the gene encoding the virulence factor internalin A (*inlA*) (27), a characteristic also shared by the Norwegian CC121 clinical isolates. Interestingly, the single *L. monocytogenes* CC121 isolated in the current study; MF7617 from soil at the edge of a university campus pond in Ås, had an intact and presumably functional copy of *inlA*. This isolate was only distantly related to the clinical CC121 isolates, separated by 195 wgMLST alleles from the nearest clinical isolate. In contrast to CC121, the other three most commonly detected CCs among the Norwegian clinical isolates – CC1, CC7 and CC8 – were relatively common also among the isolates from natural and agricultural environments, with each CC having an average prevalence of between 6% and 7.5% in the rural/urban, dairy farms, and slug isolate datasets (Figure 6A).

Since listeriosis is primarily acquired from food, the frequency distribution of CCs for *L. monocytogenes* from Norwegian food processing industry (Figure 6C) was estimated from previous work encompassing 680 isolates from five meat and four salmon processing plants, collected during 2011-2015 (47). The prevalence of lineage I isolates was <1% among the food processing industry isolates; represented by only two CC2 isolates from the meat industry. In meat processing environments, CC9 was by far the most prevalent clonal group, representing 70% of isolates. It must, however, be noted that most of the collected isolates were from two intensively sampled processing plants (34). One slug isolate and three clinical isolates (2011, 2012, 2015) but none from dairy farms or samples from rural and urban environments belonged to CC9. In salmon processing environments, CC14 was most prevalent (25%), followed by CC121 (22%), CC7 (ST7, ST732 and ST995; 18%), and CC8 (ST8 and ST551; 14%). CC14/ST14 was only represented by two clinical and two slug isolates, and not detected among isolates from rural/urban environments or dairy farms. The latter three were among the four most prevalent CCs among the Norwegian clinical isolates.

To examine the diversity of the most commonly detected *L. monocytogenes* clonal groups from Norwegian natural environments in an international context, a representative subset of reference genomes belonging to ST37, ST91, and ST451 were selected for comparative analysis using cgMLST (Figure 7). Of the >6000 examined publicly available genomes, 243 belonged to one of the three relevant STs. For ST91, a limited number of international reference sequences were available, and nearly 60% of the analysed isolates were Norwegian. This suggest that this ST could be more prevalent in Norway relative to in many other countries. For ST37, only limited clustering of Norwegian isolates relative to the international isolates was observed. In contrast, for ST91 and ST451, the Norwegian isolates appeared to cluster together with isolates from other countries, indicating that they represent internationally dispersed clones.

**Figure 7:**
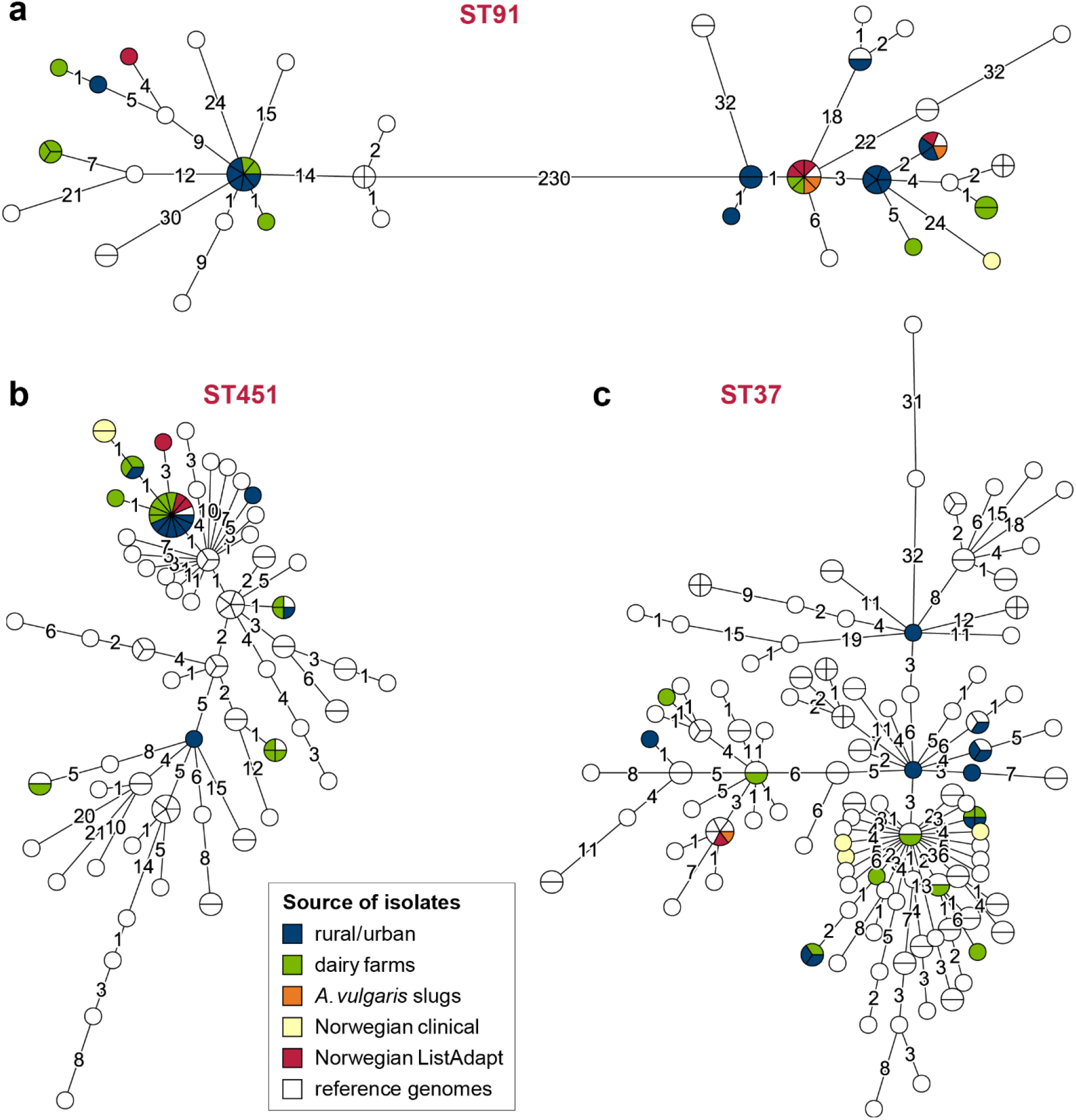
cgMLST phylogeny for the most common STs identified in the current study. Minimum spanning trees based on cgMLST allelic profiles for: **a)** ST91, **b)** ST451, and **c)** ST37, showing the relationship between the Norwegian isolates from natural environments, Norwegian clinical isolates, and reference genomes obtained from public databases. Reference genomes were obtained from the BIGSdb-*Lm* database hosted at the Pasteur Institute, WGS data from the EU project ListAdapt (also including genomes from Norwegian sources, labelled in red), and genome assemblies from NCBI GenBank. The area of each circle is proportional to the number of isolates represented, and the number of allelic differences between isolates is indicated on the edges connecting two nodes.

## Discussion

It has long been acknowledged that *L. monocytogenes* clones predominating among human clinical isolates differ from those that dominate in food (23, 24, 26, 46, 48, 49), and that persistent clones of *L. monocytogenes* may establish in food processing environments (2, 33, 34). Here, we show that the most ubiquitous clones found in soil and other natural and animal ecosystems (CC91, CC11, and CC37) are distinct from clones predominating among both clinical and food isolates, and that *L. monocytogenes* may persist and spread also in urban and rural areas, grazeland, agricultural fields, and farm environments. The correspondence of major CCs was high for the three examined sets of environmental isolates (rural/urban, dairy farms, and slugs). CC37 appeared to be exceptionally widespread in natural environments, and was isolated from nine different counties and a wide variety of habitats. It was also found to persist for years both at a farm and on a bike path in the capital of Norway. The ubiquity of this clone is also reflected by its detection in a large proportion of other studies investigating the identity of *L. monocytogenes* clones from natural and animal reservoirs, including wildlife, forest areas and farms (5, 6, 13, 18, 21, 25, 29-31, 50).

The current study identified close genetic relationships between environmental isolates of *L. monocytogenes* collected from geographically and temporally unrelated sources, despite a relatively low number of analysed isolates. Although fixed clustering thresholds for defining outbreak clusters are controversial (37, 51, 52), the genetic distances in the observed clusters were well within the thresholds used to guide outbreak analyses (23, 40-43). In the majority of observed clusters, isolates with no known likely association were indistinguishable using cgMLST analysis, which is the method currently employed for surveillance of *L. monocytogenes* by many laboratories including the Norwegian Institute of Public Health (53). This finding underscores the need for careful consideration of additional evidence such as epidemiological data, traceback evidence, and phylogenetic tree topology as part of WGS-based surveillance and outbreak investigations (54). Ideally, evaluation of possible epidemiological links should consider the occurrence of closely related strains in the whole food chain, including external contamination sources in urban and natural environments (52). Currently, a lack of published genomic data on *L. monocytogenes* from various sources is a barrier for effective management of this pathogen, both for public health authorities and for industrial actors.

During the last decade (2010-2020), an average of 24 yearly listeriosis cases have been reported in Norway, and most of them (80%) were domestically acquired (http://www.msis.no/). The implicated food is rarely identified. Only two outbreaks have been publicly reported during this period, both associated with traditional fermented fish (rakfisk); one in 2013 (ST802, 4 cases in Norway) (55), and one during the winter of 2018-2019 (ST20, 12 cases in Norway, one in Sweden (56)). A predominance of lineage II was observed among the Norwegian clinical isolates – comprising 80% of isolates during the years 2010-2015; an increase relative to the 56% observed during 1992-2005 (57). During 2010-2015, 71% of listeriosis patients were aged 70 or above, while during 1992-2005, only 42% of patients belonged to this high risk age group (http://www.msis.no/). A distinct feature among Norwegian clinical isolates was the large proportion of CC121 isolates lacking functional internalin A. The only CC121 isolate collected from a natural environment did not have an *inlA* PMSC, supporting the hypothesis that *inlA* mutations may constitute an adaptation to food industry environments (58). The relatively high proportion of cases caused by clones of a hypovirulent strain in Norway could potentially be linked to national consumption and storage practices leading to sporadic ingestion of large numbers of the pathogen among high risk groups.

Worldwide, the hypervirulent clones CC1 and CC4 are significantly more prevalent among clinical isolates than food isolates (27, 59, 60). Together, these two CCs constituted 8% of Norwegian clinical isolates and 11% of the isolates from natural and farm environments. CC1 and CC4 also appear to be prevalent in natural environments in other countries (13, 29). They were, however, not detected in a study of *L. monocytogenes* in nine Norwegian food processing plants (47). Although at least 80% of meat, cheese and fish consumed in Norway is produced domestically (61), imported processed foods remain a potential source of infections. Notably, however, 45% of Norwegian households report that they hunt, fish or collect bivalve molluscs, and about half of the population grow their own vegetables, herbs or fruit, and collect berries in the wild (62). Furthermore, the current study identified clusters comprising closely related isolates from both clinical sources and natural environments, despite comparing temporally non-overlapping sets of isolates. Together, these observations suggest that the relative contribution of industrially processed foods to listeriosis infections could be lower in Norway than in other countries.

## Materials and Methods

### Sampling of *L. monocytogenes* from rural and urban environments

Samples were taken to cover what was hypothesized as hot spots and cold spots for *L. monocytogenes* in the outer environment. The sampling plan was designed to cover different geographical regions of Norway and areas hypotesized to have respectively high (urban areas, grazeland, animal paths and areas near food processing factories) and low (forests and mountain areas, agricultural fields, beaches and sandbanks) occurrence of *L. monocytogenes*. Samples classified as footpaths were generally from non-urban areas in woods or other areas used for hiking, but separately categorized as we considered footpaths to be associated with human activities to a greater extent than more pristine woodland or mountain areas. A detailed sampling scheme was prepared and convenience sampling was performed by people living in or travelling to different areas to cover Norway geographically and to get detailed results from specific areas (e.g., gardens) and local information. The sampling was performed by trained microbiologists informed about the objective of the study and which types of sites that should be sampled. When possible, several different suspected hot and cold spots were sampled in the same geographical area, e.g., grazeland and a forest nearby where the cattle did not have access. Sampling was performed year around except for winter. For a selection of sampling sites, sampling was repeated once or more over a period of three years.

The environmental samples (soil, sand, mud, decaying vegetation, surface water, animal dung, etc.) were collected in sterile 50 mL Nunc tubes. All sampling locations were photographed, and GPS coordinates, sample content, habitat/area, and weather conditions were recorded at the time of sample collection. Specific information about the sample was also noted, such as which animals the area was frequently exposed to (e.g., cattle, deer, sheep, doves) and local information (e.g., popular area for hiking). The humidity of the collected samples was assessed on a scale from 1 to 5, ranging from completely dry (1) to liquid (5). Samples were stored at 4°C for up to a week before processing, and analysed according to ISO 11290-1 (63) with selective enrichment in half-Fraser and Fraser broth (Oxoid) and final plating on RAPID’*L*.*mono* agar (Bio-Rad).

### Whole genome sequencing

For each *L. monocytogenes* isolate from rural/urban environments or from *Arion vulgaris* slugs (21), a single colony was picked, inoculated in 5 mL brain heart infusion broth, and grown at 37°C overnight. Culture samples (1 mL) were lysed using Lysing Matrix B and a FastPrep instrument (both MP Biomedicals) and genomic DNA was isolated using the DNeasy Blood and Tissue Kit (Qiagen). Libraries for genome sequencing were prepared using the Nextera XT DNA Sample Preparation Kit (Illumina) and sequenced using 2×300 bp reads on a MiSeq instrument (Illumina).

Colonies from the *L. monocytogenes* isolates from dairy farms (16) were inoculated in 20 mL tryptone soy broth and incubated at 37°C for 24 h before 1 mL was pelleted and DNA extracted using the DNeasy Blood & Tissue kit (Qiagen). Libraries for genome sequencing were prepared using the NEBNext Ultra DNA Library Prep Kit (New England Biolabs) with random fragmentation to 350 bp and sequencing of 2×150 bp on a NovaSeq 6000 S4 flow cell (Illumina).

### Genome assembly

All genome assemblies used in phylogenetic analysis were generated as follows: Raw reads were filtered on q15 and trimmed of adaptors before *de novo* genome assembly was performed using SPAdes v3.10.0 or v3.13.0 (64) with the careful option and six k-mer sizes (21, 33, 55, 77, 99, and 127). Contigs with sizes of <500 bp and with coverage of <5 were filtered out. For the *L. monocytogenes* isolates from dairy farms, the genomes released to NCBI GenBank as PRJNA744724 <u>(see Data availability section) were generated</u> using SPAdes v3.14.1, incorporated in the software tool Shovill available at https://github.com/tseemann/shovill. Shovill also performed adaptor trimming using Trimmomatic, corrected assembly errors, and removed contigs with sizes <500 bp and coverage <2. The quality of all assemblies was evaluated using QUAST v5.0.2 (65) (results in S9 Table).

### Phylogenetic analyses

Classical MLST analysis followed the MLST scheme described by Ragon *et al*. (66) and the database maintained at the Institute Pasteur’s *L. monocytogenes* online MLST repository (https://bigsdb.pasteur.fr/listeria/). *In silico* MLST typing was performed for raw sequencing data using the program available at https://bitbucket.org/genomicepidemiology/mlst (67), and for genome assemblies using the program available at https://github.com/tseemann/mlst. CCs are defined as groups of ST profiles sharing at least six of seven genes with at least one other member of the group, except for CC14, which is divided into CC14 represented by ST14 and ST399 in the current work, and CC91 represented by ST91, as isolates belonging to these two groups do not cluster together in phylogenetic analyses of *L. monocytogenes* populations (27).

The wgMLST analysis was performed using a whole-genome scheme containing 4797 coding loci from the *L. monocytogenes* pan-genome and the assembly-based BLAST approach, implemented in BioNumerics 7.6 (https://www.bionumerics.com/news/listeria-monocytogenes-whole-genome-sequence-typing). The cgMLST analysis was performed using the scheme described by Moura *et al*. (23), which is a subscheme of the wgMLST scheme employed in the BioNumerics platform. For publicly available genomes (see below), cgMLST profiles were obtained by sequence query against the BIGSdb-*Lm* cgMLST allele database maintained at the Institut Pasteur (https://bigsdb.pasteur.fr/listeria/). For the genomes sequenced in the current study, cgMLST profiles were extracted from the wgMLST profiles by mapping of the sequences of the cgMLST allele subset to the publicly available nomenclature, through synchronization of BioNumerics with the BIGSdb-*Lm* cgMLST allele database. A subset of isolates were subjected to cgMLST analysis using both approaches to confirm that identical cgMLST profiles were obtained. During wgMLST analysis in BioNumerics, each identified unique allele sequence is designated an allele identifier integer. In contrast, for analyses involving the BIGSdb-*Lm* cgMLST allele database, only alleles which are already present in the database will be identified and receive an allele identifier, while novel alleles are recorded as missing loci.

Minimum spanning trees were constructed using BioNumerics based on the categorical differences in the allelic cgMLST or wgMLST profiles for each isolate. The number of allelic differences between isolates was read from genetic distance matrices computed from the absolute number of categorical differences between genomes. Loci with no allele calls were not considered in the pairwise comparison between two genomes. The criterion for inclusion of a cluster in S2 Table, S3 Table, S4 Table, S5 Table, and S7 Table was that each genome included in the cluster showed ≤20 or ≤21 wgMLST allelic differences towards at least one other genome in the cluster. For S6 Table, clusters comprising isolates showing ≤10 cgMLST allelic differences towards at least one other genome in the cluster were included. Consequently, for clusters with three or more genomes, individual pairs of genomes with genetic distances exceeding the set thresholds can and will be included in the clusters (see also S1 Text).

### Publicly available genomes

Available genomes of clinical isolates from human patients in Norway were identified by searching the NCBI Pathogen Detection database (https://www.ncbi.nlm.nih.gov/pathogens) on the 30.08.2021. Available raw sequencing data from NCBI BioProjects submitted by ECDC and NPIH; PRJEB26061 (45) and PRJEB25848, were subjected to *de novo* genome assembly as described for isolates from rural/urban environments. *In silico* MLST genotyping was successful for all genomes except one of the genomes published by the ECDC, and wgMLST analysis was successful for all except 21 of the ECDC genomes.

Reference genomes included in the cgMLST analysis of ST37, ST91 and ST451 genomes were identified from the following selected sources on the 27.08.2021: (i) cgMLST profiles from the BIGSdb-*Lm* database (https://bigsdb.pasteur.fr/listeria/); 15 genomes belonging to relevant STs were identified. (ii) Raw WGS data from the ListAdapt project (https://onehealthejp.eu/jrp-listadapt/) comprising 1552 genomes (BioProject PRJEB38828). *De novo* genome assembly was performed for the 165 genomes of relevant STs. (iii) Genome assemblies from NCBI GenBank; among the 3926 *L. monocytogenes* genomes, 63 genomes belonged to the relvant STs.

## Data availability

Data from this whole-genome shotgun project has been deposited at the NCBI GenBank database under BioProjects PRJNA689486, PRJNA744724, and PRJNA689487. For GenBank and Sequence Read Archive (SRA) accession numbers, see S1 Table. The assemblies were annotated using the NCBI Prokaryotic Genomes Automatic Annotation Pipeline (PGAAP) server (http://www.ncbi.nlm.nih.gov/genome/annotation_prok/).

## Acknowledgments

The authors wish to thank Merete Rusås Jensen, Anette Wold Åsli, Janina Berg, Tove Maugesten at Nofima for excellent technical assistance. We also sincerely thank collaborators in the food and salmon processing industry, colleagues at Nofima (Runar Gjerp Solstad, Rasmus Karstad, Halvor Nygaard, Kristin Skei Nerdal) and Arild Hugo Solstad for contributing to sample collection. We thank Ann-Katrin Llarena (Faculty of Veterinary Medicine, Norwegian University of Life Sciences) for help during genome assembly of *L. monocytogenes* dairy farm isolates. We thank Ida Skaar (Norwegian Veterinary Institute) for providing the *L. monocytogenes* isolates from slugs (21). We thank the team of curators of the Institut Pasteur MLST system (Paris, France) for importing novel alleles, profiles and/or isolates at http://bigsdb.pasteur.fr/listeria/. This work was funded by the Norwegian Agriculture and Food Industry Research Funds, grant numbers 262306 and 314743.

## List of supplemental information

**S1 Text** (*Supplemental_S1_Text*.*docx*): Additional information regarding clusters of closely related *L. monocytogenes* isolates.

Supplemental Tables (*Supplemental_Tables*.*xlsx*):

**S1 Table**: Isolates included in the current study

**S2 Table**: Clusters of isolates from rural or urban environments

**S3 Table**: Clusters of dairy farms isolates

**S4 Table**: Clusters containing isolates from both rural/urban environments and dairy farms

**S5 Table**: Clusters containing isolates from slugs or both slugs and rural/urban/farm environments

**S6 Table**: Clusters of clinical isolates

**S7 Table**: Clusters containing both clinical isolates and isolates from rural/urban/farm environments or slugs

**S8 Table**: Survey of STs and CCs found in previous studies of various natural and farm environments

**S9 Table:** Results from evaluation of genome assemblies using QUAST

