## Supplemental Text S1 for "WGS analysis of *Listeria monocytogenes* from rural, urban, and farm environments in Norway: Genetic diversity, persistence, and relation to clinical and food isolates"

### Supplemental Material
### S1 Text

Additional information regarding clusters of closely related *L. monocytogenes* isolates.

**WGS analysis of *Listeria monocytogenes* from rural, urban, and farm environments in Norway: Genetic diversity, persistence, and relation to clinical and food isolates**

Annette Fagerlund[a]#, Lene Idland[b], Even Heir[a], Trond Møretrø[a], Marina Aspholm[b], Toril Lindbäck[b], and Solveig Langsrud[a]

[a] Nofima, Norwegian Institute of Food, Fisheries and Aquaculture Research, Ås, Norway

[b] Department of Paraclinical Sciences, Faculty of Veterinary Medicine, Norwegian University of Life Sciences, Ås, Norway

#### Persistent strains detected in rural and urban environments

Persistence events in rural and urban environments were revealed by WGS analysis (S2 Table). Sampling in a private garden compost heap on three occasions resulted in seven positive samples of *L. monocytogenes*. Four isolates identified in 2016 and 2019 belonged to ST451 (CC11), and three ST425 (CC90) isolates were from 2019 and 2020. The ST451 isolates differed by 1 to 10 wgMLST allelic differences, with 2 to 9 alleles distinguishing the 2019 isolate from the three isolates from 2016. In the ST425 cluster, two isolates from 2020 were indistinguishable by wgMLST but differed by 2 alleles relative to the strain isolated 4 months earlier, in 2019.

Sampling in the town centre of Ås and the nearby NMBU university campus (Akershus) in September 2019 and January 2020 also resulted in detection of several clones that were repeatedly collected from the same sampling site. One was a ST398 (CC19) clone, first isolated in front of a park bench by a flower bed in 2019. In 2020, an isolate from the same location and an isolate from the adjacent flower bed were found, with all three isolates differing from each other by 2 wgMLST alleles. Two ST20 (CC20) isolates, indistinguishable by wgMLST, were collected in 2019 and 2020, from two sampling sites by the university campus pond, located only 5 meters apart. Two isolates belonging to ST204 (CC204), separated by only 3 allelic differences, were collected two years apart (2017 and 2019) from the same flowerbed next to the entrance to a subway station in central Oslo. Sampling in Oslo also resulted in detection of two ST37 (CC37) isolates differing by 7 alleles, isolated 4 months apart from samples of sand/gravel on a bike path next to a major road.

In some cases, isolates belonging to the same clone were collected from nearby locations, but not at the exact same sampling site. For example, sampling in rural areas in the vicinity of Ås resulted in detection of a cluster of two pairs of ST91 (CC91) isolates collected 4 months apart, which were separated by 2 to 5 wgMLST allelic differences. One of the isolates was found ~500 meters north of the area where the remaining three were found, but both sites were used as pasture for sheep. It is, therefore, possible that the same herd of sheep may have grazed and shed *L. monocytogenes* at both locations. Interestingly, two years earlier, a related isolate differentiated from this cluster by 11 to 12 allelic differences was collected from a flower bed in Ås town centre, around 2 km to the southwest. Furthermore, two isolates, belonging to a different clade within ST91 and separated by only 8 allelic differences, were collected in 2016 and 2017 at sampling sites separated by 1.5 km in the Ås area.

Sampling in a port town in western Norway, in an area with operational fish processing industry, resulted in isolation of several clones of *L. monocytogenes* which were identified during multiple occasions. During a period of almost 1 year, two isolates belonging to ST1 (CC1) differing by only 4 wgMLST alleles, and eight isolates belonging to ST732 (CC7), differing by 0 to 23 wgMLST alleles, were collected at in different locations on the quay. The source of these isolates is likely to be the fishing industry, since the same clones were also identified within the fish processing plant located in the same area (data not shown). However, four isolates belonging to ST647 (CC20), which had not been detected inside the factory, were also isolated from the same area. These four isolates, two from 2018 and two from 2020, differed by 2 to 3 alleles.

#### Persistence and cross-contamination on Norwegian dairy farms

Potential cross-contamination events on the dairy farms, e.g., contamination of milk (assessed by sampling milk filters) from feed or feces found on the same farm, were revealed by WGS analysis (S3 Table). Notably, only three out of 12 isolates from milk filters were closely related (1-3 wgMLST allelic differences) to fecal and/or feed isolates obtained during the same sampling occasion at the same farm (farms 1 and 12). In all three cases, the same clone (max 5 wgMLST allelic differences) was isolated from feed samples also during subsequent sampling occasions. Furthermore, the same ST91 clone – identical by wgMLST analysis – was collected from a milk filter and a teat swab during the same sampling occasion (farm 13). A ST451 clone found on a milk filter on the second visit to farm 12 was found to be closely related (6-7 allelic differences) to isolates from feed and feces samples collected during a later farm visit. We also observed a case where two ST451 isolates from a milk filter and from feed collected during the same visit to farm 9 were less likely to be from the same contamination source, as the two isolates differed by 28 wgMLST alleles. Nevertheless, it seems clear that milk filters (and consequently milk) may become contaminated with *L. monocytogenes* clones found in the farm environment.

Out of the 19 visits to a farm where both the feed and feces samples were positive for *L. monocytogenes*, the same clone (0-3 wgMLST allelic differences) was found in both samples on seven occasions. An additional four cases were identified where the same clone (0-3 allelic differences) was found on a milk filter or teat swab and in feed or feces samples, or in both milk filter and teat swab samples, on the same visit. Six of these clones were also isolated during more than one visit to the same farm. All in all, we found 12 pairs or clusters of isolates that were repeatedly isolated from the same farm, with pairwise allelic differences within each cluster ranging from 0 to 11 wgMLST alleles. The interval between visits ranged from 2 to 10 months. Four of the clusters were found on Farm 12, which was the farm with the highest detection rate for *L. monocytogenes* with 15 positive samples in total. The remaining eight repeatedly isolated clones were collected on eight different farms. Together, these findings strongly suggest that the same *L. monocytogenes* clones can persist over time in dairy farm environments. The identified persistent clones belonged to ST4, ST8, ST18 and ST2761 (both CC18), ST37, ST91, ST177, ST226, ST394 (CC415), ST412, and ST451 (CC11; two clusters).

#### Detection of closely related isolates from different geographic areas

Four cases where the same clone was collected on more than one farm were identified (S3 Table). In the first case, two identical ST394 (CC415) isolates collected from feed samples from Farm 2 in January and February 2020 showed 11 wgMLST allelic differences towards an isolate collected from feces at Farm 5 in November 2019. The feed sample taken on the same visit at Farm 5 was positive for ST226, not ST394. In the second case, this ST226 isolate from feed from Farm 5 showed 9 wgMLST allelic differences towards a feed sample from the same farm isolated three months earlier, and 12 allelic differences towards an isolate from feed at Farm 4 also obtained in November 2019. The three farms were located in the same geographical area of Oppland county. In the third case, involving ST451 (CC11) and Farms 8 and 9, located in Akershus county, a milk filter isolate from one farm showed 19 allelic differences towards a feed sample from the second farm. The fourth case also involved ST451 and a total of eight strains; a cluster of three isolates from feed and feces from Farm 6, and five isolates obtained from milk filter and feces samples on four other farms. The number of allelic differences between these isolates (not considering the differences within the Farm 6 cluster) ranged from 15 to 57, with isolates from all farms linked by 20 or fewer pairwise allelic differences. The farms were not located in the same geographical area (Østfold, Akershus and Oppland). These data suggest that farms located at different geographical areas may host the same genetic clones of *L. monocytogenes*. Although the diversity between clones found on different farms was somewhat greater than the diversity between clones found on the same farm, the isolates from different farms could in most cases not be distinguished using the commonly employed core genome MLST (cgMLST) analysis.

A total of six clusters were identified containing *L. monocytogenes* isolates from both rural or urban environments and dairy farms, with genetic distances ranging from 9 to 27 wgMLST allelic differences, and 0 or 1 cgMLST differences (S4 Table). Several of the links involved isolates obtained from livestock grazeland in the Ås area. The closest link was observed between two ST37 isolates collected in the vicinity of Ås. These samples were collected 3 years and 1.5 km apart and differed by only 3 alleles. Both locations were grazing land/pasture, with one of the locations also used as a feeding location for livestock. These two isolates were closely related to two isolates from feed and teat swab samples obtained on two different visits to farm 12, about 50 km from Ås, with 9-14 wgMLST allelic differences between the pairs of isolates. In another ST37 cluster, an isolate from feed from dairy farm 1, located north of Oslo, showed 15 and 16 wgMLST allelic differences towards two linked isolates from grazing land/pasture from an area in southwestern Norway. Similarly, the previously described ST91 cluster, consisting of four isolates collected in the period 2019-2020 from grazing land close to Ås, showed 20 or 21 allelic differences compared to an isolate from feed at farm 18, located about 100 km north-west of Ås. Furthermore, an ST6 isolate collected from a feed sample from dairy farm 17, located about 70 km west of Ås, differed by 14 wgMLST alleles from an isolate collected from grazing land at Ås in 2020. These two isolates were part of a cluster also containing an isolate collected next to a tree in central Oslo in 2017, and which differed from the two other isolates with 18 and 9 alleles, respectively. Two additional cases where dairy farm isolates differed from strains from the rural/urban area dataset with about 20 allelic differences were also found. The first was the cluster of three ST394 isolates from Farms 2 and 5, which showed 21 or 23 wgMLST allelic differences towards an isolate from farmland in northern Norway from 2018. The second involved the cluster of eight ST451 strains from five different farms, which differed by between 15 and 27 wgMLST alleles from the cluster of ST451 isolates from the garden compost heap in Ås.

#### Genetic distances between clinical and environmental isolates

Nine clusters containing both one or more clinical isolates and one or more isolates originating from rural and urban environments, dairy farms, or slugs is presented in S7 Table. One cluster belonged to CC8 (ST8) and two belonged to CC7 (ST7). The ST8 cluster comprised a slug isolate from 2012 with 14-15 allelic differences towards two clinical isolates from 2012 and 2013. Within ST7, the first cluster contained three clinical isolates from 2010 and 2015 differing by 10 to 17 wgMLST alleles. This cluster also contained an isolate from slugs with 9 to 18 allelic differences towards the three clinical isolates, and an isolate taken in the vicinity of a horse paddock in Oslo in 2020, showing 12 to 21 differences towards the clinical isolate trio. The other ST7 cluster was composed of five closely related clinical isolates, one from 2010 and the remaining four from 2012, separated by only 1 to 6 allelic differences. This cluster was genetically associated, through allelic differences ranging from 11 to 15, with a single isolate from 2020 obtained from a sample taken in the woods in the vicinity of a meatpacking factory. Strikingly, there was another relatively good match between an isolate taken in the vicinity of this factory and a clinical isolate. These were ST220 isolates linked by 19 wgMLST allelic differences, one obtained in 2020 from the road leading to the factory and the other a clinical isolate from 2013. The two Norwegian clinical CC11 (ST451) isolates – both from 2013 and separated by 2 allelic differences – were linked by 9 and 11 wgMLST allelic differences to an isolate from 2019 obtained from a milk filter on a dairy farm. These three isolates were part of a larger cluster of relatively closely related ST451 isolates – with seven additional dairy farm isolates and six isolates from rural or urban locations separated from the two clinical isolates by distances ranging from 14 to 23 wgMLST alleles.

#### Regarding differences in genetic distances between cgMLST and wgMLST analyses

The cgMLST scheme (1748 loci) is a subscheme of the wgMLST scheme (4797 loci in total). In S6 Table, clusters of clinical isolates showing of ≤10 cgMLST allelic differences towards at least one other genome in the cluster were included. When the genomes in each of these clusters were analysed using wgMLST, the genetic distances within these clusters ranged from 2 to 105. In S2-S4

Tables and S7 Table, clusters of genomes showing maximum 20-21 wgMLST allelic differences towards at least one other genome in the cluster were included. Also in these clusters, there were several examples of clusters with few cgMLST allelic differences and a relatively large range of variable genes in the wgMLST analysis.

Two factors contributed to the large difference in variable genes obtained using cgMLST analysis relative to wgMLST analysis observed in a subset of the clusters. Both factors were associated with the fact that during pairwise comparison between two genomes, a locus is not called as variable if one of the genomes does not have an allele call for this locus.

Firstly, several cgMLST loci in which the alleles differed between genomes were not recorded as such because the allele found in one or more of the genomes in a cluster was not present in the BIGSdb-*Lm* cgMLST allele database. This effect could be alleviated by a greater representation of genomes in the BIGSdb-*Lm* cgMLST isolate database.

Secondly, the cgMLST scheme containing core loci per definition does not contain variable genetic elements, while the wgMLST scheme contains stable loci from the accessory genome, including loci in prophage regions. If one genome in a cluster lacks a certain prophage, these loci will (correctly) not be called in the wgMLST analysis. However, if two or more other genomes do show allelic variations in these prophage genes, the range of pairwise wgMLST distances between genomes in the cluster can become relatively large. Each genome containing the prophage is nevertheless linked to the cluster by a small number of genetic differences towards the genome(s) lacking the prophage(s) in question.

Examples:

The two CC177 genomes in **S6 Table Cluster 13** (ERR2522308, ERR2522327) differed by 73 wgMLST alleles but only 2 cgMLST alleles. For 32 of the differences called by wgMLST, the locus was part of the cgMLST subscheme but only called in one genome in the pair in the cgMLST analysis. Of the 39 differing wgMLST loci that were not part of the cgMLST subscheme, 17 were located in regions identified as prophage sequences using the PHASTER phage search tool (https://phaster.ca/).

**Cluster 24 in S3 Table** contains eight genomes belonging to CC11/ST451. The genetic distance within the cluster (not considering the differences between the three closely related isolates D118L, D117L, D044L from Farm 6) ranged from 0-1 cgMLST alleles and 15-57 wgMLST alleles. In total, 105 wgMLST loci showed variable alleles in at least one pair of genomes, and of these, 36 loci belonged to the cgMLST subscheme. Overall, 34 of the differing cgMLST loci were not reported as variable in the cgMLST analysis because the least frequent allelic variant was not present in the BIGSdb-*Lm* cgMLST allele database. Of the 69 variable loci not present in the cgMLST subscheme, 44 were located in regions identified as prophage sequences using the PHASTER phage search tool, and one locus belonged to a plasmid.

The eight CC7 genomes in **S6 Table Cluster 23** differed by 0-10 cgMLST alleles and by 1-105 wgMLST alleles. A total of 176 wgMLST loci were called as variable among the genomes in the cluster, of which 86 belonged to the cgMLST subscheme. Of these, 76 were not reported as variable in the cgMLST analysis because the least frequent allelic variant was not present in the BIGSdb-*Lm* cgMLST allele database. Of the 90 variable loci not present in the cgMLST subscheme, 39 were located in regions identified as prophage sequences using the PHASTER phage search tool, and one locus belonged to a plasmid.
